# Distributed and diverse hindbrain neuronal activity contributes to sensory processing and motor control in the *Xenopus laevis* tadpole

**DOI:** 10.1101/2021.06.10.447865

**Authors:** Giulia Messa, Stella Koutsikou

## Abstract

Locomotion is a key feature of healthy animals, which depends on their ability to move -or not to move-for their survival. The hatchling *Xenopus laevis* tadpole responds to trunk skin stimulation by swimming away, and its developing nervous system is simple enough to make it an ideal model organism to study the control of locomotion. This vertebrate embryo relies on excitatory cells in the skin to detect the sensory stimulus, which is quickly sent to the brain via ascending sensory pathway neurons. When the stimulation is strong enough, descending reticulospinal neurons are activated in the hindbrain and the spinal cord, after a long and variable delay. The activation of reticulospinal neurons indicates the initiation of swimming and sustains the rhythmic firing of CPG (central pattern generator) neurons, among which are motor neurons. The tadpole is then able to rhythmically contract trunk muscles, allowing the undulatory movement of swimming. However, how the tadpole’s developing brain exerts descending control over reticulospinal neurons, and thus over the spinal CPG centers, is not fully understood yet. In this work, we recorded extracellular activity in the hindbrain of the tadpole to identify firing units that are involved in the long and variable delay to swim initiation following trunk skin stimulation. We isolated firing units that mediate distinct motor output. We subsequently grouped them in populations based on their firing patterns in response to skin stimulation and motor output. We propose a novel neural circuitry for sensory processing and descending motor control exerted by the hindbrain of the hatchling tadpole, which could account for the long and variable delay to reticulospinal neuron activation, and thus swim initiation.

## Introduction

In nature, animals must select, execute and adapt their motor behavior to specific aims, such as feeding, escaping from a predator or finding a mate, with the ultimate goal of surviving and reproducing (1). From nudibranchs (2) to lamprey (3), from cats (4) to humans (5), all animals are required to coordinate and timely activate central and peripheral neuronal circuits, in order to perform the most advantageous movement. Most of the sensory input animals receive from the environment leads to motor activity. This can be driven by the nature of the stimulus (*e.g.* withdrawal from the source of harmful heat), or by the instinctive need to inspect the surroundings (*e.g.* eye movement to follow a moving object).

In adult vertebrates, the combined activation of neural circuits in the forebrain, brainstem and spinal cord generates a fine-tuned and rich repertoire of motor behaviours, including locomotion (1, 6). Spinal CPG (central pattern generator) circuits are activated and via a sophisticated integration of ipsilateral activation and contralateral inhibition of CPG neurons (7, 8) enable the animal to move efficiently (9, 10). Yet, and although the spinal neural circuits have been extensively studied in higher vertebrates (11, 12), the cellular underpinnings of some critical features of mammalian locomotion have not been completely unraveled (for example the flexor-extensor coordination; (13)).

On the contrary, the spinal neuronal circuits responsible for swimming in the *Xenopus laevis* tadpole have been extensively characterized, and the reticulospinal interneurons (namely descending interneurons, dINs) that drive spinal CPG activity have been anatomically and functionally described (14–16). Therefore, the simplicity and detailed knowledge of the tadpole’s neuronal network that drives and sustains locomotion makes this simple animal an ideal model organism to study the supraspinal mechanisms of descending motor control.

Different brain regions have been identified as key components for the generation of movement (13), with the basic neural circuit being conserved from lamprey to humans (13). The basal ganglia play a major role in descending control over the MLR (mesencephalic locomotor region) (6,7). This circuit relies on tonic inhibition of the MLR at resting conditions, provided by GABAergic neurons from GPi (globus pallidus pars interna) and SNr (substantia nigra pars reticulata) (2). When striatal projections silence these highly active inhibitory neurons, the MLR excitation over reticulospinal neurons (RS) is released (8,9). Recently, key glutamatergic neurons in the MLR have been found to be sufficient to induce locomotion in mice (17) and, in a more caudal region of the mouse brainstem, the LPGi (lateral paragigantocellular nucleus), the activation of vGlut2-expressing neurons has been shown to be sufficient to start locomotion and increase its speed (10).

However, unravelling the supraspinal control of movement in higher vertebrates (18–20) has been challenging, and the neuron-to-neuron pathways that coordinate even the simplest motor decision and movement execution, have not been fully elucidated (21–25).

It has been shown that the hatchling *Xenopus laevis* tadpole responds to trunk skin stimuli by initiating swimming with a delay of 50-150ms (26–28). We demonstrated that this long and variable delay resembles the accumulation of excitation proposed for complex brain circuits active during motor decision-making (29, 30), in contrast to quick reflexes (*e.g* the quick C-start response; (31)), thus concluding that sensory processing and motor decision mechanisms are present even at this early developmental stage. In particular, whole-cell recordings of hindbrain reticulospinal neurons (dINs) showed synaptic potentials that contribute to a slow and variable increase in excitation, which in turn allowed us to infer the existence of candidate pre-synaptic neurons, namely hindbrain extension neurons (hexNs; (26)). Ultimately, and only if the stimulus delivered is strong enough, this build-up of excitation in the dINs reaches a firing threshold, which in turn marks the start of swimming (16, 28).

The sensory (Rohon-Beard cells, RB) and sensory pathway neurons (dorsolateral ascending and dorsolateral commissural neurons, dla and dlc, respectively), which transmit the sensory information from the periphery to the tadpole’s brain, are very well-known (19,20). Their firing, early in response to stimulation, cannot explain neither the variability, nor the delay detected in dINs’ firing latencies and swim initiation (27). Indeed, the variable and long process of accumulation of excitation recorded in dINs could be mimicked through modelling only by inserting an excitatory recurrent network within the brainstem sensory pathway (26).

Despite the evidence described above, which indicate that the fundamental elements of descending motor control are present at an early developmental stage, the neurons constituting the proposed neuronal network in the *Xenopus* tadpole hindbrain are currently undefined. In this work, we implemented a top-down approach to uncover the firing characteristics of the neuronal components responsible for the accumulation of excitation in reticulospinal interneurons, prior to movement execution. Specifically, we recorded extracellular activity in the hindbrain of the tadpole, and we analyzed single unit firing patterns during distinct motor states, including swim initiation.

We identified novel firing patterns across the tadpole’s hindbrain that indicate an overall increased neuronal activity in the time between the sensory stimulation and motor output. We discriminated four new neuronal subpopulations based on their firing activity, which differentially contributed to accumulation of excitation in the hindbrain prior to the start of swimming. We discuss how these newly identified neuronal populations are accountable for the motor decision-making process (to swim or not to swim) taking place in the simple embryonic brain of the *Xenopus laevis* tadpole. Importantly, we also propose a simple neural network that includes the newly identified neuronal populations and their connections within the existing well-defined sensory, sensory-pathway and CPG neuronal circuitry in the tadpole’s hindbrain and spinal cord.

## Results

Extracellular hindbrain neuronal activity was recorded in response to stimulation, and subsequently analysed in relation to fictive swimming in the *Xenopus laevis* tadpole. Electrical trunk skin stimulation was delivered at intensities above and below the threshold for swimming, and hindbrain neuronal activity was studied in four distinct motor states: 1) rest, 2) stimulation which was not followed by swim initiation (stim/no start), 3) stimulation which led to swim initiation (stim/start), and 4) sustained swimming (swimming) (fig. 1Aiv). Spike sorting analysis (see Materials and Methods) was used to isolate and subsequently categorise single units. All units were mapped along the rostro-caudal axis of the tadpole’s hindbrain, which was arbitrarily divided into three sectors as shown in fig. 1Aiii.

**Figure 1.**
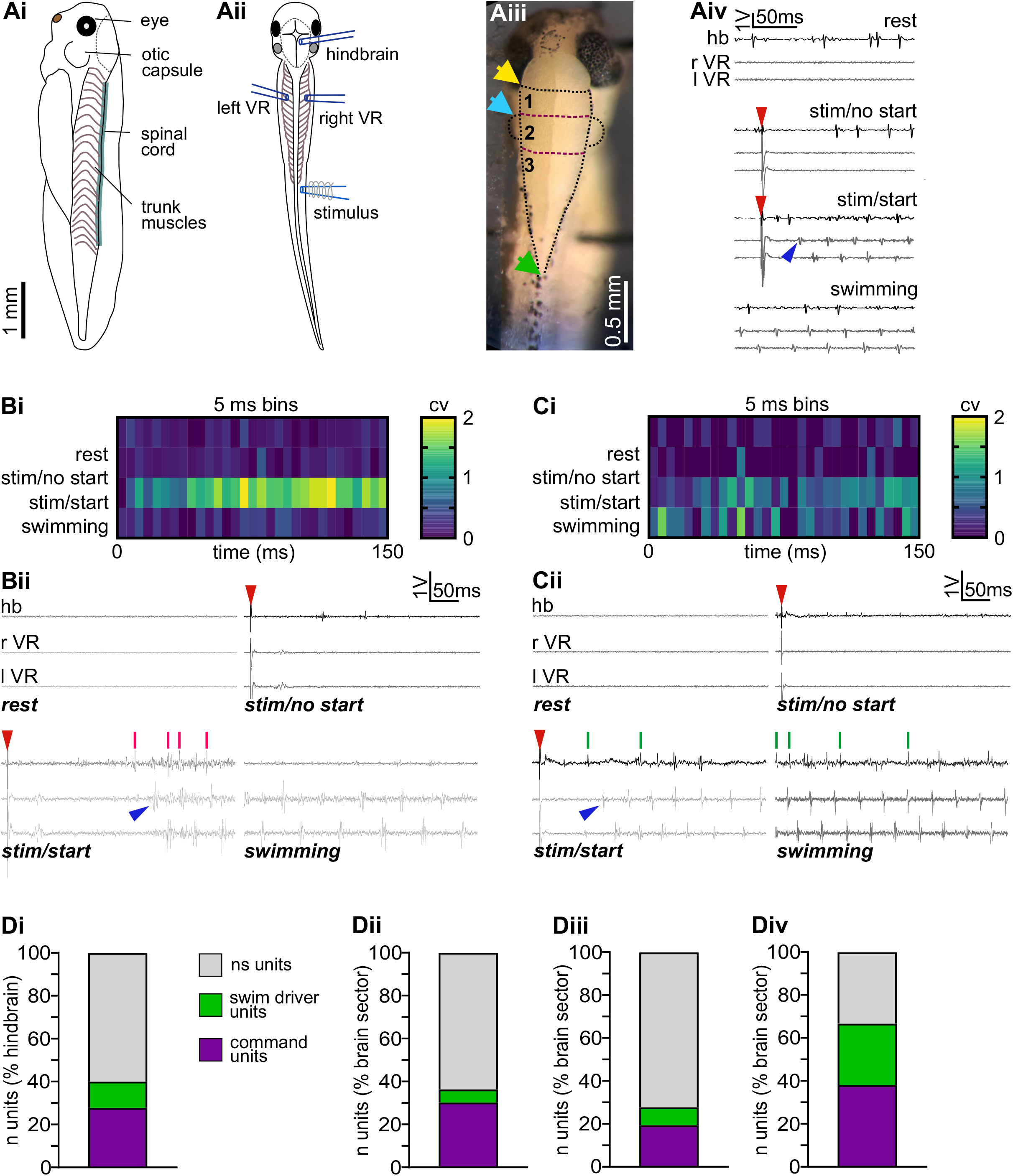
A) Experimental design. Ai. Lateral view of stage 37/38 *Xenopus laevis* tadpole; anatomical features are indicated on the head (eye and otic capsule) and along the body (spinal cord and trunk muscles). Grey dotted line on the head represents the area where the skin was removed to expose the brain.
Aii. Dorsal view of stage 37/38 *X. laevis* tadpole with extracellular suction electrodes positioned as per experimental conditions. Grey dotted line marks the area where the skin was removed to make the hindbrain accessible. In different experiments, the brain electrode (hindbrain) was positioned in one of the three hindbrain sectors as depicted in fig. 1Aiv. The two VR (ventral root) electrodes were positioned at the 5^th^ inter-myotomal cleft on both sides of the tadpole’s body (left VR, right VR). The stimulating electrode (stimulus) was attached to the skin on the right side of the body at the level of the anus. Scale bar as in Ai.
Aiii. *X. laevis* tadpole’s hindbrain as it appears when the skin is removed. The hindbrain was visually divided into three sectors along the rostro-caudal axis using well-defined anatomical landmarks, *i.e.* MHB (yellow arrow), otic capsules (blue arrow) and obex (green arrow): rostral sector (1), otic capsule level (2) and caudal sector (3).
Aiv. Examples of the four different motor states analysed (200 ms): rest, stimulation/no start of swimming, stimulation/start of swimming, sustained swimming. Raw traces for hindbrain extracellular activity (hb, black trace) and right and left VR recording (rVR and lVR, grey traces) are presented. Red arrowheads indicate stimulus delivery, blue arrowhead indicates the start of swimming. **B, C) command units and swim driver units are activated at the start of swimming**. Bi) Average heat map of command units recorded in the hindbrain (25 units in 11 animals, minimum of 4 trials per motor state per units). Coefficients of variation (CV=standard deviation/mean) were calculated for each time bin (5 ms) on the number of spikes fired by single units during the same 5 ms in the four motor states.
Bii) Examples of spikes fired by one command unit recorded in the four motor states investigated (rest, stimulation/no start, stimulation/start, swimming) 150 ms are reported for each example. Red lines indicate spikes fired by the unit, and they are presented above the respective extracellular hindbrain recording raw trace (black trace). Fictive swimming is shown by rhythmic right and left VR bursts (grey trace, rVR and lVR). Red arrowheads represent the delivery of the electrical stimulus; blue arrowhead indicates the start of swimming.
Ci) Average heat map of swim driver units recorded in the hindbrain (11 units in 8 animals, minimum of 4 trials per motor state per units). Coefficients of variation (CV=standard deviation/mean) were calculated for each time bin (5 ms) on the number of spikes fired by single units during the same 5 ms in the four motor states. **Cii)** Examples of spikes fired by one swim driver unit recorded in the four motor states investigated (rest, stimulation/no start, stimulation/start, swimming) 150 ms are reported for each example. Green lines indicate spikes fired by the unit, and they are presented above the respective extracellular hindbrain recording raw trace (black trace). Fictive swimming is shown by rhythmic right and left VR bursts (grey trace, rVR and lVR). Red arrowheads represent the delivery of the electrical stimulus; blue arrowhead indicates the start of swimming. **D) Distribution of command and swim driver units** Di. Bar charts representing percentages for non-significant (grey), swim driver (green) and command units (violet) in the entire hindbrain (Di), and in the different hindbrain sectors as depicted in Aiv (**Dii**, rostral sector; **Diii** otic capsule level; **Div**, caudal sector). Percentages were calculated over the total number of units recorded in the hindbrain (Di, 54/90 = 60.0% non significant units; 11/90 = 12.2% swim driver units, 25/90 = 27.8% command units), and over the total number of units recorded in each of the hindbrain sectors (Dii-rostral sector, 21/33 = 63.6% non significant units, 2/33 = 6.1% swim driver units, 10/33 = 30.3% command units; Diii-otic capsule level, 26/36 = 72.2% non significant units, 3/36 = 8.3% swim driver units, 7/36 = 19.4% command units; Div-caudal sector, 7/21 = 33.3% non significant units; 6/21 = 28.6% swim driver units, 8/21 = 38.1% command units;).

### Novel hindbrain firing patterns involved in control of swimming and its initiation

We identified two novel firing patterns that were correlated with swim initiation in the tadpole. Specifically, we detected 25 units out of 90 (recorded in 18 animals), which showed significantly higher activity at the start of swimming in response to skin stimulation (‘stimulus X swim interaction’ units, *p*<0.05, two-way ANOVA; 150 ms were analysed starting from stimulation, fig. 1Bi, Bii). We named these units, with firing rate higher only at motor initiation, ‘command units’. We also detected 11 units (out of 90 units in 18 animals) that showed increased firing activity both at motor initiation and during sustained swimming (‘swim effect’ units, *p*<0.05, two-way ANOVA; fig. 1Ci, Cii). These units, which were more active both at the start of movement, as well as during continuous swimming, were named ‘swim driver units’. Command units were highly active only when the electrical stimulus was strong enough to induce swimming in the tadpole (the experimental condition referred to as ‘stimulation/start’), while they were mainly silent at rest, during sustained swimming, and when the stimulation failed to trigger swimming (fig. 1B). On the other hand, swim driver units showed increased firing activity both at swim initiation caused by the stimulus delivered to the trunk skin, as well as during sustained swimming. These units were mostly inactive when the animal was not swimming, *i.e.* at rest and when stimulation did not lead to swimming (fig. 1C).

Both command and swim driver units were recorded throughout the hindbrain (fig. 1D), with an overall prevalence for command units over swim driver units (27.8% for command vs 12.2% for swim driver units, fig. 1Di). In the rostral area of the hindbrain (sector 1, fig. 1Aiii), 30.3% of the recorded units were command units (10/33 units), while 6.1% (2/33 units) fell into the swim driver category (fig. 1Dii). At the level of the otic capsule (sector 2, Fig. 1 Aiii), 19.4% of the units were found to be command units (7/36 units) and 8.3% were swim driver units (3/36 units) (fig. 1Diii). The highest percentage of command and swim driver units was recorded in the most caudal area of the hindbrain (sector 3, Fig. 1Aiii), where 38.1% of the units recorded were command units (8/21 units), and 28.6% (6/21 units) were swim driver units. (fig. 1Div). Percentages are calculated over the total number of units recorded within a single hindbrain sector.

All other units recorded (54 out of 90) did not change their firing rate in response to trunk skin stimulation, or to swimming behaviour (‘non-significant units’, *p*>0.05, two-way ANOVA). These units showed stable, low firing in every motor state tested (rest, stimulation/no start, stimulation/start and swimming). Therefore, we concluded that these units were not implicated in the control of swim initiation. Such units were recorded throughout the hindbrain: 63.6% in the rostral area (21/33 sector’s units), 72.2% (26/36 sector’s units) at the otic capsule level, 33.3% (7/21 sector’s units) in the caudal area (fig. 1Dii–iv, percentages calculated over the total number of units recorded within each hindbrain sector). These units were excluded from subsequent analysis.

Furthermore, CPG neurons are also activated as soon as swimming starts. None of the swim driver units studied in this work showed features of a CPG-like activity.

### Command units are activated earlier than swim driver units, and both firing patterns are distributed throughout the hindbrain

Command unit firing started increasing prior to swim initiation, leading to a peak of activity (0.48 ± 0.21 Hz, mean ± SEM) 11 ms before the start of locomotion (fig. 2Ai). After the movement had started, command units slowly decreased their firing rate, till they became silent during sustained swimming (fig. 2Ai). Swim driver units also increased their activity rate before the start of swimming (fig. 2Ai), but a peak in their firing activity was not detected. Instead, swim driver unit firing rates were stable as swimming became continuous (0.06 ± 0.08 Hz 10 ms before start vs 0.06 ± 0.09 Hz 10 ms after swimming initiation; data expressed as mean ± SEM). Spikes fired by command units were recorded earlier before movement initiation in comparison to the spikes fired by swim driver units (fig. 2Aii).

**Figure 2.**
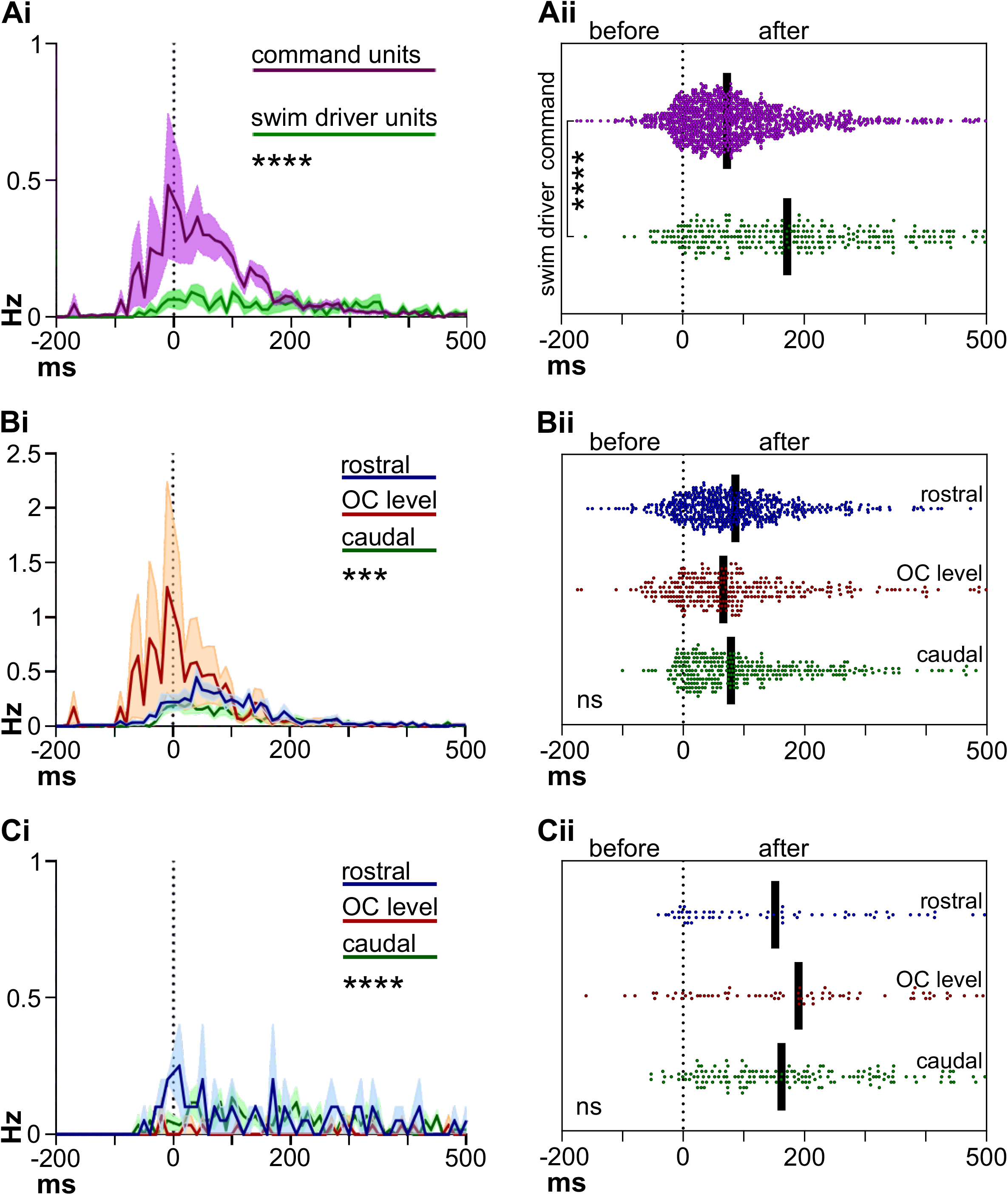
A) Command and swim driver units are activated at different latencies. Ai) Firing rates of command (violet line) and swim driver units (green line) recorded before movement initiation and in the first 500 ms of sustained swimming. Dotted grey line (ms=0) marks the start of fictive swimming. Data are presented as mean (solid lines) ± SEM (shaded area). Wilcoxon matched pairs signed rank test, *p*<0.0001; command units 0.0925 ± 0.0139, swim driver units 0.0299 ± 0.0031 (mean ± SEM). Command units N=25, swim driver units N=11; minimum 4 trials/unit. \*\*\*\**p*<0.0001
Aii) Scatter plot of spikes fired by command units (violet) and swim driver units (green) before movement initiation and during the first 500 ms of sustained swimming. Dotted grey line (ms=0) marks the start of fictive swimming. Black solid lines indicate median values (median ± SD= 72.69 ± 104.4 ms for command units; median ± SD = 160.8 ± 144.3 ms for swim driver units). Kolmogorov-Smirnov test, *p*<0.0001; 1145 spikes from 25 command units, 288 spikes from 11 swim driver units. \*\*\*\**p*<0.0001 **B, C) Command units and swim driver units show different firing rates throughout the hindbrain** Bi) Firing rates of command units recorded in the rostral sector (blue), at the otic capsule level (red) and in the caudal sector (green) of the hindbrain. Dotted grey line (ms=0) marks the start of fictive swimming. Data are presented as mean (solid lines) ± SEM (shaded area). Repeated measures one-way ANOVA with Bonferroni correction for multiple comparisons, *p*=0.0003; rostral vs OC level, mean diff.=0.0725; rostral vs caudal, mean diff.= 0.0376; OC level vs caudal, mean diff.= 0.1101. Rostral sector N=10 units; otic capsule level N=7 units; caudal sector N=8 units; minimum 4 trials per unit. \*\*\**p*<0.001
Bii) Scatter plot of spikes fired by command units recorded in the rostral sector (blue), at the otic capsule level (red) and in the caudal sector (green) of the hindbrain. Dotted grey line (ms=0) marks the start of fictive swimming. Black solid lines indicate median values (median ± SD = 71.94 ± 93.29 ms for rostral units; median ± SD = 65.97 ± 117.5 ms for otic capsule level units; median ± SD= 78.32 ± 109.5 ms for caudal units). Kruskal-Wallis test, *p*=0.0618; 558 spikes from 10 rostral units, 284 spikes from 7 otic capsule level units, 303 spikes from 8 caudal units.
Ci) Firing rates of swim driver units recorded in the rostral sector (blue), at the otic capsule level (red) and in the caudal sector (green) of the hindbrain. Dotted grey line (ms=0) marks the start of fictive swimming. Data are presented as mean (solid lines) ± SEM (shaded area). Repeated measures one-way ANOVA with Bonferroni correction for multiple comparisons, *p*<0.0001; rostral vs OC level, mean diff.= 0.0390; rostral vs caudal, mean diff.= 0.0137; OC level vs caudal, mean diff.= −0.0254. Rostral sector N=2 units; otic capsule level N=3 units; caudal sector N=6 units; minimum 4 trials per unit. \*\*\*\**p*<0.0001
Cii) Scatter plot of spikes fired by swim driver units recorded in the rostral sector (blue), at the otic capsule level (red) and in the caudal sector (green) of the hindbrain. Dotted grey line (ms=0) marks the start of fictive swimming. Black solid lines indicate median values (median ± SD = 127.8 ± 148.6 ms for rostral units; median ± SD = 190.2 ± 160.7 ms for otic capsule level units; median ± SD = 162.2 ± 134.1 ms for caudal units). Kruskal-Wallis test, *p*=0.2837; 63 spikes from 2 rostral units, 73 spikes from 3 oc level units, 152 spikes from 6 caudal units.

Command units recorded in the central area of the hindbrain (otic capsule level) were responsible for the firing rate’s peak at swimming initiation, with a peak of 1.28 ± 2.23 Hz recorded 10 ms before the start (fig. 2Bi). Command units detected in the rostral and caudal sectors increased their firing in a less abrupt fashion (fig. 2Bi), contributing nevertheless to the overall augmented activity, which was prolonged after the initiation of swimming (0 to ~150 ms after the start, fig. 2Ai). However, the onset and distribution of the spikes fired at the start of movement by command units in the three hindbrain sectors did not differ (fig. 2Bii).

Swim driver units recorded in the three hindbrain areas showed prolonged firing rate patterns at the initiation of swimming (fig. 2Ci), resulting in a constant activity that persisted as swimming became sustained. The distribution of the spikes fired by swim driver units did not differ among the three hindbrain areas (fig. 2Cii).

Both command and swim driver units showed a distribution of activity throughout the three hindbrain sectors, leading us to consider the two populations to be highly dispersed throughout the hindbrain.

### Subpopulations of command and swim driver units are differentially activated

Based on their distinctive firing patterns, we identified two subgroups among command units. The first group, which we named ‘first level command units’, showed increased firing in the stimulation/start trials (fig. 3Ai and Aii), but was also slightly active when the stimulus delivered was not strong enough to cause swim initiation (fig. 3Bi and Bii). On the contrary, the second group, named ‘second level command units’, was active only when the stimulation led to motor response (fig. 3Ai, Aii and 3Bi, Bii). In the time prior to swim initiation, spikes fired by first level command units were recorded at shorter latencies from swim initiation than those fired by second level command units (fig. 3C, negative area of the graph). After swimming had started, the temporal distribution of spike fired by first and second level command units did not differ (fig. 3C, positive area of the graph).

**Figure 3.**
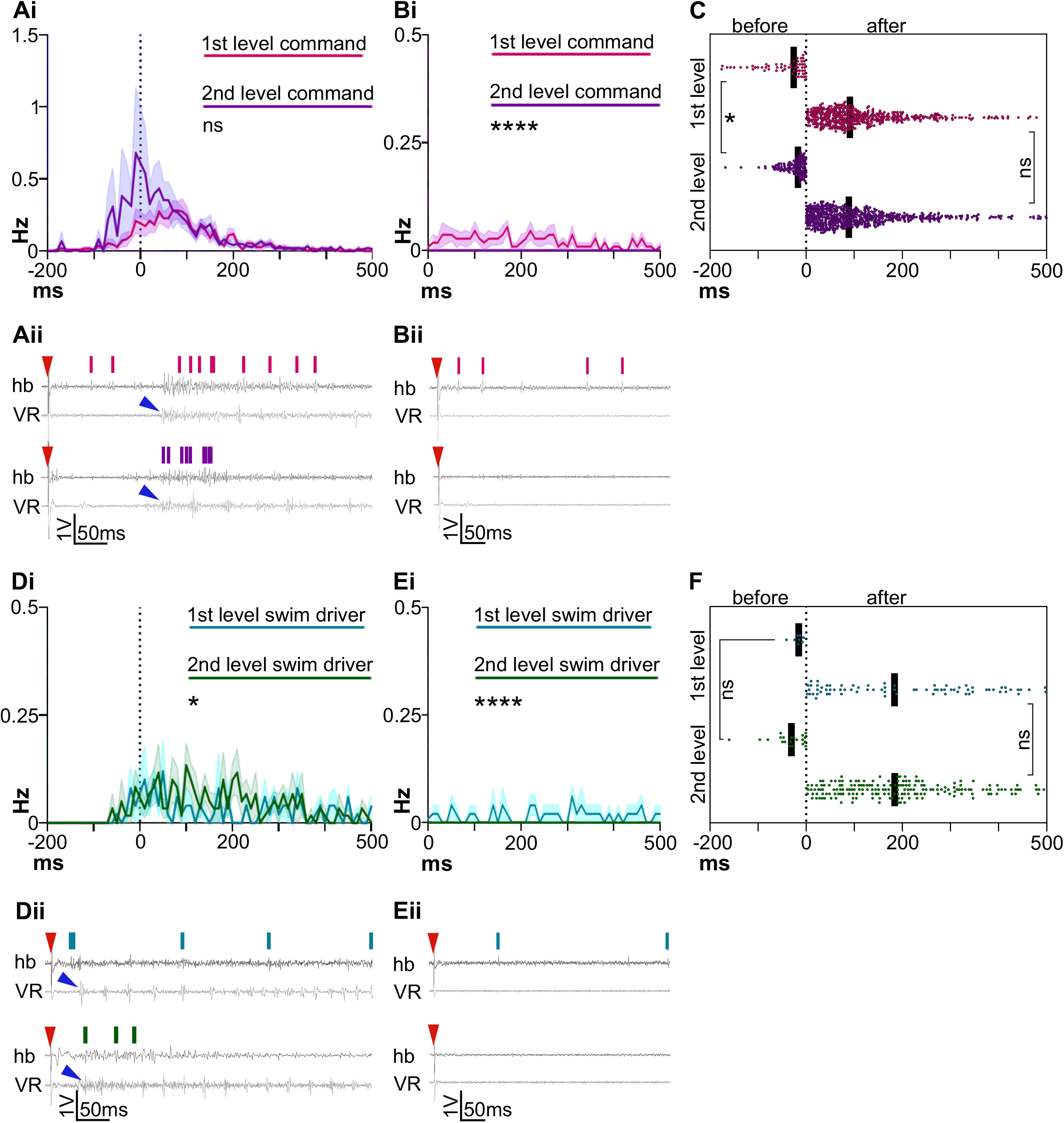
Subpopulations of command and swim drive units are activated differently. Ai) Firing rates of first (pink line) and second level (violet line) command units recorded before movement initiation and in the first 500 ms of sustained swimming. Dotted grey line (ms=0) marks the start of fictive swimming. Data are presented as mean (solid lines) ± SEM (shaded area). Wilcoxon matched pairs signed rank test, *p*=0.0511; first level command units 0.0695 ± 0.0100 ms, second level command units 0.1094 ± 0.0180 ms (mean ± SEM). First level command units N=11, second level command units N=14; minimum 4 trials/unit.
Aii) Examples of spikes fired by one first level command unit (top trace, pink lines) and one second level command units (bottom trace, violet lines), recorded in the stimulation/start motor state. Spikes fired by the units are presented above the respective extracellular hindbrain recording raw trace (hb, black trace). Fictive swimming is shown by rhythmic VR bursts (grey trace, VR). For clarity, only the VR with the first burst, marking swimming initiation, is shown here. Red arrowheads represent the delivery of the electrical stimulus; blue arrowhead indicates the start of swimming.
Bi) Firing rates of first level (pink line) and second level (violet line) command units recorded in the first 500 ms after stimulation. Data are presented as mean (solid lines) ± SEM (shaded area). Wilcoxon matched pairs signed rank test, *p*<0.0001; first level command units 0.0149 ± 0.0018 ms, second level command units 0.0000 ± 0.0000 (mean ± SEM). First level command units N=11, second level command units N=14; minimum 4 trials/unit. \*\*\*\**p*<0.0001
Bii) Examples of spikes fired by one first level command unit (top trace, pink lines) and one second level command unit (bottom trace, no lines), recorded in the in the stimulation/no start motor state. Spikes fired by the units are presented above the respective extracellular hindbrain recording raw trace (hb, black trace). The absence of fictive swimming is shown by the silent VR (grey trace, VR). As both VR were silent in this case, for clarity only one VR is shown here. Red arrowheads represent the delivery of the electrical stimulus.
C) Scatter plot of spikes fired by first level command units (pink) and second level command units (violet) before movement initiation and during the first 500 ms of sustained swimming. Dotted grey line (ms=0) marks the start of fictive swimming. Black solid lines indicate median values. Kolmogorov-Smirnov test on data recorded before swim initiation, *p*=0.0187; first level command units −25.75 ± 50.27 ms; second level command units ™17.19 ± 26.90 ms (median ± SD). 60 spikes from 11 first level command units, 120 spikes from 14 second level command units. Kolmogorov-Smirnov test on data recorded after swim initiation, *p*=0.0923; first level command units 91.09 ± 94.18 ms; second level command units 88.95 ± 104.5 ms (median ± SD). 450 spikes from 11 first level command units, 565 spikes from 14 second level command units. \**p<0.05*
Di) Firing rates of first level (blue line) and second level (green line) swim driver units recorded before movement initiation and in the first 500 ms of sustained swimming. Dotted grey line (ms=0) marks the start of fictive swimming. Data are presented as mean (solid lines) ± SEM (shaded area). Wilcoxon matched pairs signed rank test, *p*<0.0394; first level swim driver units 0.0237 ± .0035, second level swim driver units 0.0350 ± 0.0042 (mean ± SEM). First level swim driver units N=5, second level swim driver units N=6; minimum 4 trials/unit. \**p*<0.05
Dii) Examples of spikes fired by one first level swim driver unit (top trace, blue lines) and one second level swim driver units (bottom trace, green lines), recorded in the in the stimulation/start motor state. Spikes fired by the units are presented above the respective extracellular hindbrain recording raw trace (hb, black trace). Fictive swimming is shown by rhythmic VR bursts (grey trace, VR). For clarity, only the VR with the first burst, marking swimming initiation, is shown here. Red arrowheads represent the delivery of the electrical stimulus; blue arrowhead indicates the start of swimming.
Ei) Firing rates of first level (blue line) and second level (green line) swim driver units recorded in the first 500 ms after stimulation. Data are presented as mean (solid lines) ± SEM (shaded area). Wilcoxon matched pairs signed rank test, *p*<0.0001; first level swim driver units 0.0111 ± 0.0017, second level swim driver units 0.0000 ± 0.0000 (mean ± SEM). First level swim driver units N=5, second level swim driver units N=6; minimum 4 trials/unit. \*\*\*\**p*<0.0001
Eii) Examples of spikes fired by one first level (top trace, blue lines) and one second level swim driver unit (bottom trace, no lines), recorded in the in the stimulation/no start motor state. Spikes fired by the units are presented above the respective extracellular hindbrain recording raw trace (hb, black trace). The absence of fictive swimming is shown by the silent VR (grey trace, VR). For clarity, only one VR is shown here. Red arrowheads represent the delivery of the electrical stimulus.
F) Scatter plot of spikes fired by first level (blue) and second level swim driver units (green) before movement initiation and during the first 500 ms of sustained swimming. Dotted grey line (ms=0) marks the start of fictive swimming. Black solid lines indicate median values. Kolmogorov-Smirnov test on data recorded before swim initiation, *p*=0.1182; first level command units −15.65± 11.20 ms; second level swim driver units −30.61 ± 38.49 ms (median ± SD). 8 spikes from 5 first level swim driver units, 21 spikes from 6 second level swim driver units. Kolmogorov-Smirnov test on data recorded after swim initiation, *p*=0.0803; first level swim driver units 183.5 ± 152.7 ms; second level swim driver units 183.9 ± 125.3ms (median ± SD). 75 spikes from 5 first level swim driver units, 184 spikes from 6 second level swim driver units.

Similarly, the swim driver population could be divided into two subpopulations. First level swim driver units were active when stimulation was delivered to the animal, irrespective of the motor outcome (fig. 3Di, Dii and 3Ei, Eii). On the other hand, second level swim driver firing was detected only when the electrical stimulus led to swimming (fig. 3Di, Dii and 3Ei, Eii). Contrary to command units, both first and second level swim driver units showed the same distribution of spikes prior to movement initiation (fig.3F, negative area of the graph), as well as after swimming had become continuous (fig. 3F, positive area of the graph).

## Discussion

This work presents the first evidence of the distributed and diverse hindbrain neuronal excitability accounting for the long and variable latency to *X. laevis* tadpole swim initiation. Using threshold and subthreshold trunk skin electrical stimuli evoking distinct motor outputs, we categorised hindbrain units based on their firing patterns and latencies in relation to the initiation of swimming. We identified two highly distinct groups of the previously so-called hindbrain extension neurons (hexNs; (26)), and based on their firing properties, we named them ‘command’ and ‘swim driver’ units.

We showed that both hexN types had the ability to extend the sensory memory based on their variable firing latency and frequency, followingr stimulation above and below the threshold for swimming (fig. 2A). Their firing patterns are also in agreement with well-established theories on sensory memory and motor decision-making, based on the existence of a variable accumulation of excitation to a threshold for movement initiation (30, 32–34).

Furthermore, these units’ firing patterns cannot be ascribed to any of the well-known cell types of the tadpole central nervous system (15). The firing of both types of units differed significantly from the early and mostly single-spike firing of sensory pathway neurons (dla and dlc) (35), as well as the rhythmic and late firing of dINs, key in the initiation and maintenance of locomotor patterns (16, 26). However, our data showed that both command and swim driver units could act pre-synaptically to dINs. Indeed, both types of units fired earlier than the start of locomotion indicated by the first VR burst, and with variable latencies across trials. This is in agreement with both the latency of synaptic potentials previously recorded on dINs, as well as their long and variable firing (26).

Furthermore, we identified subtypes of both command and swim driver units. Both groups’ subtypes were categorised as ‘first level’ and ‘second level’ units based on their firing patterns in response to subthreshold stimuli, which did not lead to initiation of fictive swimming. Indeed, first level units across both groups fired in response to subthreshold stimuli, even if at lower frequency, when compared to the firing after suprathreshold stimulation. This is in full agreement with the presence of synaptic potentials and accumulation of excitation on dINs (26) even when the stimulus does not lead to dIN firing and initiation of fictive swimming.

Based on our current findings, we propose a supraspinal mechanism of descending motor control, which includes the newly identified hindbrain units, as depicted in fig. 4. We suggest that command units work as sensory processors in the hindbrain of the tadpole, being postsynaptic to sensory pathway neurons (dla and dlc), whilst swim driver units act at later stages, providing the necessary excitation to dINs in the hindbrain. This is supported by swim driver’s average spiking latency, which is longer compared to command units’ firing. Initially, the sensory information received by Rohon-Beard cells in the skin is carried to the brain by dlc and dla neurons, it is then weighted and integrated by the proposed sensory processing centre in the hindbrain, comprised by first and second level command units (fig. 4). When the stimulus delivered is strong enough to lead to movement, first and second level command units fire and will excite both subpopulations of swim driver units (stimulation/start; fig. 4A), which will in turn provide the cumulative excitation to dINs, allowing them to reach their firing threshold. The firing of dINs will lead to motor neuron excitation and the initiation of the undulatory movement of swimming (16, 36).

**Figure 4.**
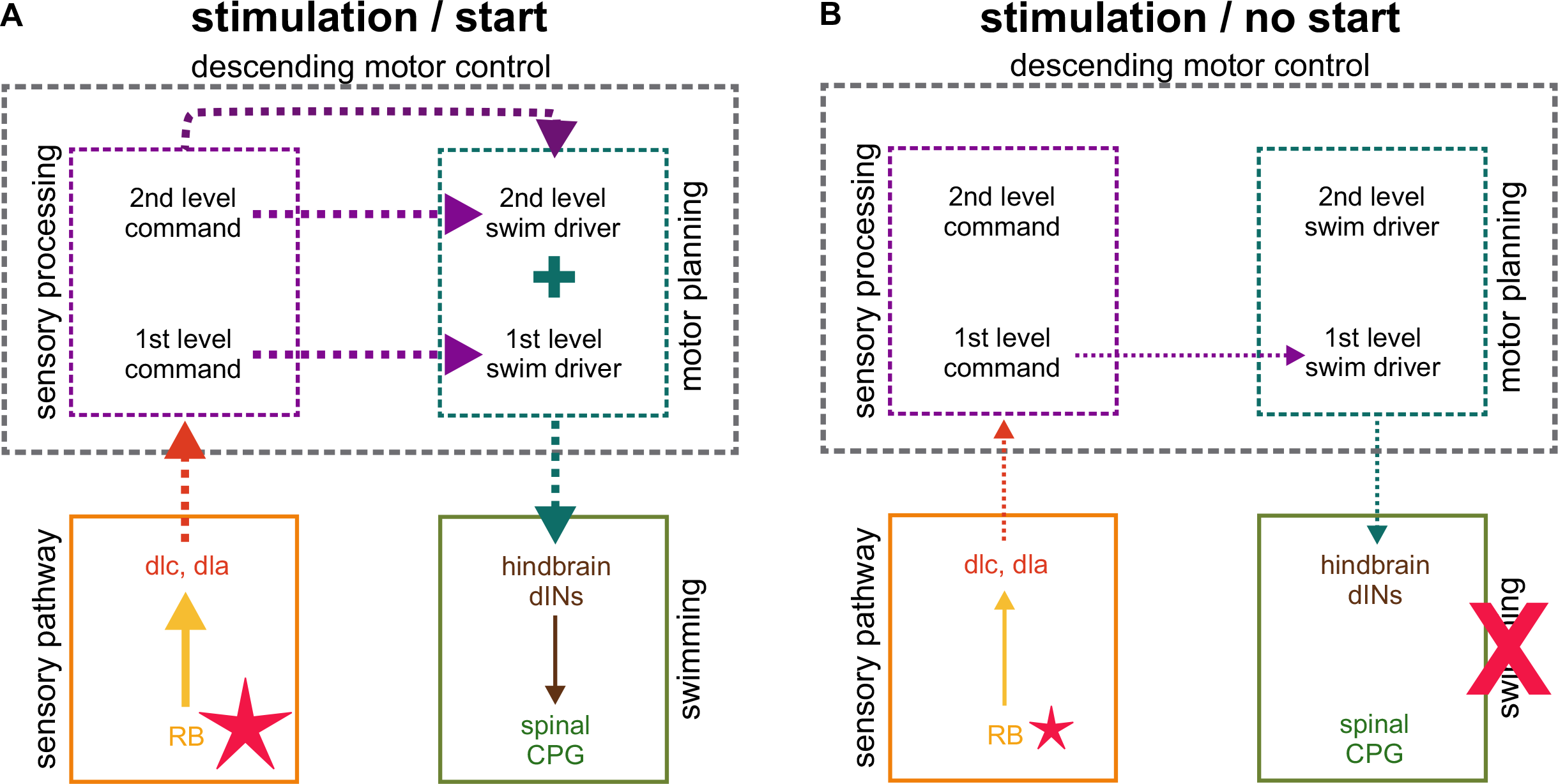
Proposed neural mechanism for sensory processing and motor descending control. A) Scheme of the proposed neural circuit active when a suprathreshold stimulus is delivered to the tadpole’s trunk skin leading to swim initiation (stimulation/start). Solid arrows represent known synaptic connections, solid line boxes indicate known circuits (sensory pathway and swimming). Dotted arrows and boxes represent proposed connections and circuits in the hindbrain (sensory processing, motor planning, descending motor control). A higher firing rate is represented by thicker arrows, compared to the same arrows in B. Red star represents stimulation that is strong enough to lead to swim initiation.
B) Scheme of the proposed neural circuit active when a subthreshold stimulus is delivered to the tadpole’s trunk skin, failing to initiate swimming (stimulation/no start). Solid arrows represent known synaptic connections, solid line boxes indicate known circuits (sensory pathway and swimming). Dotted arrows and boxes represent proposed connections and circuits in the hindbrain (sensory processing, motor planning, descending motor control). A lower firing rate in the various synaptic connections is represented by thinner arrows, compared to the same arrows in A. Smaller than in A, red star represents a weaker stimulation, which does not lead to swim initiation. Red ‘X’ indicates that the tadpole does not start to swim.

All first level units were active also when the trunk skin stimulus applied was below the threshold for swimming (stimulation/no start, fig. 4B), in contrast to second level units which were inactive in the same condition. The firing of first level units will still lead to depolarisation of dINs, but only below their firing threshold, thus not allowing swim initiation.

Furthermore, we hypothesise that the second level command population is less likely to fire due to its electrical membrane properties providing the neuronal circuit with the means to discriminate between stimulus intensities. In this scenario, second level command units, as well as first level ones, will receive synaptic input from dlc and dla neurons, however they will not be activated due to their higher firing threshold. On the contrary, first level command units will be activated at lower stimulus intensities. Once active, second level command units would excite second level swim driver units, which will provide, together with first level swim drivers, strong excitation to dINs (fig. 4A). A different firing likelihood for first and second level command populations might also explain the slightly delayed firing of second level units before swimming starts, compared to first level command units (fig. 3C).

Although it is not possible to precisely locate the neuronal somata through extracellular recordings, we discovered that swim driver unit firing was preferentially localised in the caudal portion of the hindbrain (fig. 1Div), while command units firing was more dispersed along the hindbrain. This anatomical layout might partially reflect the function of the distinct neuronal populations, *i.e.* swim driver units would be excited, and thus controlled, by command units. This layout across the tadpole hindbrain is in agreement with studies in complex vertebrate brains, where neurons involved in motor decision-making and planning possess diverse spatial and temporal firing profiles, and they are intermingled across different brain areas (37).

In this study we provide the first direct evidence of the spatial and temporal ‘extension’ of sensory information across the tadpole’s hindbrain. We attribute to this hindbrain neuronal activity a major role in the accumulation of excitation on reticulospinal neurons, whose firing, or lack of, will in turn manifest into the tadpole’s binary motor decision to swim or not to swim, respectively. We believe that the identification of the neuron-to-neuron pathway(s) and how individual cells modulate aspects of the tadpole’s behavior are the important next steps in unravelling the role of supraspinal brainstem control on motor output.

## Materials and Methods

### Ethics, animal care and preparation

*Xenopus laevis* embryos were supplied by the European *Xenopus* Resource Centre (EXRC; Portsmouth, UK). Animal care and all experimental procedures on *Xenopus laevis* tadpoles were approved by the University of Kent Animal Welfare and Ethical Review Body (AWERB) committee. *Xenopus laevis* tadpoles at developmental stage 37/38 (38) were used and all experiments were conducted at room temperature (19-22°C).

Tadpoles were briefly anesthetized with 0.1% MS-222 (ethyl 3-aminobenzoate methanesulfonate, Sigma-Aldrich), and subsequently immobilized by immersion in a 10 μM α-bungarotoxin (Invitrogen) solution for 50 minutes. Both solutions mentioned above were made in saline (NaCl 115 mM, HEPES 10 mM, NaHCO3 2.4 mM, KCl 3 mM, MgCl2 1mM, CaCl2 2mM) adjusted to pH 7.4. Animals were then mounted onto a rotating Sylgard block submerged in saline, and dissection was carried out as previously described (14, 28). Briefly, the skin covering the brain and trunk muscles on both sides was removed, giving access to the entire hindbrain and to myotomal clefts (14, 28). The trigeminal nerves were severed at both sides of the body to prevent initiation of swimming in response to propagation of skin impulse (39–41).

### Electrophysiology

Extracellular ventral root recordings, indicative of fictive swimming, in combination with extracellular recordings of hindbrain neuronal activity, were performed on immobilized *X. laevis* tadpoles at stage 37/38. Two borosilicate glass suction electrodes (tip diameter ~50μm) filled with saline were attached to both sides of the tadpole’s body (fig.1Ai, Aii), approximately at the level of the 5^th^ myotomal cleft (14, 28). A third glass suction electrode (tip opening ~30μm) filled with saline was used to record extracellular hindbrain neuronal activity (fig.1Ai, Aii). The hindbrain recording electrode was randomly positioned in one of the three hindbrain areas depicted in fig. 1Aiii. The electrode’s location was annotated based on its position relative to anatomical landmarks, *i.e.* the midbrain-hindbrain border (MHB), otic capsules and the obex (fig. 1Aiii). Ventral root and hindbrain extracellular activity were amplified, filtered and digitized via a 1041Power (CED, Cambridge, UK) and recorded in Signal 7 (CED, Cambridge, UK). Electrical stimulation was delivered in single square pulses through a glass suction electrode, wrapped in silver wire and filled with saline. This stimulating electrode was attached to the trunk skin at the level of the anus (fig. 1Aii). Both intensity (V) and duration (ms) of the stimulus were set in each experiment as the smallest values required to evoke fictive swimming. All animals initiated fictive swimming after a stimulation within the range of 3.5-4.5 V and 0.25-0.4 ms. Ventral root and hindbrain neuronal activity were recorded in four motor states: 1) at rest, when no stimulus was applied to the tadpole’s skin and ventral root activity was absent; 2) stimulation/no start, when the stimulus delivered was not strong enough to produce fictive swimming; 3) stimulation/start, when the stimulus delivered triggered fictive swimming; 4) swimming, during sustained fictive swimming (fig. 1Aiv).

### Data analyses

Spike sorting, based on single spikes’ size and shape, was carried out on all hindbrain extracellular recordings using Spike2 10.00 (CED, Cambridge, UK), and single units were visually evaluated for spike shape consistency. The number of spikes fired by individual units was counted during 150 ms trials in each of the four motor states (fig. 1Aiv). Randomly chosen 150 ms repetitions throughout the recording were analysed for ‘rest’ and ‘swimming’ states. For the two states where stimulation was applied (‘stimulation/start’ and ‘stimulation/no start’), the time frame analysed was stimulation (t=0) +150 ms. A minimum number of 4 trials were analysed for each of the four motor states in each experiment. Two-way ANOVA with Geisser-Greenhouse correction was run (GraphPad Prism 9) on the number of spikes counted for each unit in the four motor states described above. Stimulation and swimming were the two factors tested in the two-way ANOVA. Depending on the statistical outcome of the two-way ANOVA, units considered for further classification and firing pattern analysis were: 1) units which showed a *p* value <0.05 for the interaction between stimulation and swimming; 2) units which showed a *p* value <0.05 for the swimming factor (swim effect). Coefficient of variations (CV=standard deviation/mean) were used to create the heat maps presented in the figures. CV was calculated for each time bin (5 ms) on the number of spikes fired by each unit during the same 5 ms.

## Acknowledgements

The authors would like to acknowledge the support of The Physiological Society UK through a Research Grant awarded to SK and The University of Kent through a PhD Scholarship awarded to GM.

